# A *pbpB1* mutation causing β-lactam resistance in clinical *Listeria monocytogenes* isolates

**DOI:** 10.1101/2025.04.22.649998

**Authors:** Sabrina Wamp, Rosalyn Wagner, Franziska Schuler, Alexander Krüttgen, Antje Flieger, Sven Halbedel

## Abstract

**Objectives:** Listeriosis is a severe foodborne infection and associated with high mortality. Treatment is based on ampicillin, amoxicillin or penicillin, often combined with gentamicin, but meropenem is also used occasionally. β-lactam resistant *Listeria monocytogenes* isolates are infrequently described but the mechanism of resistance is not known.

**Methods:** A clinical *L. monocytogenes* isolate with suspected β-lactam resistance was collected from a German listeriosis patient. Resistance profiling, whole genome sequencing, comparative genomics and genetic experiments were used to identify the causative DNA polymorphism. Spontaneous ampicillin resistant suppressors of *L. monocytogenes* reference strain EGD-e were selected and their genomes sequenced.

**Results:** A W428R substitution near the active site of penicillin binding protein B1 (PBP B1) was identified as the cause of ampicillin, amoxicillin and meropenem resistance. Further clinical *L. monocytogenes* isolates with similar resistance profiles were found by searching the genome database of the German consultant laboratory for *Listeria* for *pbpB1 W428R*-positive isolates. Furthermore, the same mutation was selected for in strain EGD-e during cultivation in the presence of ampicillin.

**Conclusions:** *L. monocytogenes* can develop β-lactam resistance by a specific substitution in PBP B1, likely selected for during β-lactam treatment. Antibiotic susceptibility testing should be considered an important part of adequate listeriosis therapy.

## Introduction

Listeriosis has one of the highest case fatality rates among the foodborne bacterial infections (up to ∼30%) (1). The disease is caused by the bacterium *Listeria monocytogenes*, a non-sporulating, rod-shaped bacterium belonging to the phylum *Bacillota* that is omnipresent in many environmental reservoirs, from where it is transmitted to humans through contaminated food (2). Episodes of asymptomatic carriage in the gut is quite common (3), but as listeriosis is usually self-limiting in healthy individuals, the incidence of diagnosed infections is low. However, elderly patients, pregnant women and neonates as well as individuals with immunocompromising disease or those receiving immunosuppressive therapy or proton pump inhibitors are at risk to develop invasive infections (4). During invasive listeriosis, the bacteria translocate from the gut to the blood stream, then disseminate to the liver and the spleen and may later infect the central nervous system or the placenta of pregnant women (2). During pregnancy-associated listeriosis, horizontal transmission either occurs through *in utero* infection of the foetus after hematogenous spread of the bacterium to the placenta or during birth due to vaginal colonization originating from the gastrointestinal tract (5). In adult patients, invasive listeriosis manifests as meningitis, rhombencephalitis, sepsis or fever. Pregnancy associated listeriosis is usually mild for the mother but can have adverse health effects for foetuses and neonates and may include miscarriage, stillbirth or neonatal infections such as granulomatosis infantiseptica (6).

Antibiotic therapy of listeriosis is mainly based on the administration of penicillin G or aminopenicillins (ampicillin or amoxicillin), that are often combined with gentamicin to increase their effectiveness (7). The primary lethal target of (amino)penicillins in *L. monocytogenes* is penicillin binding protein B1 (PBP B1, previously: PBP3) (8, 9). PBP B1 is a transpeptidase that catalyses the crosslinking of the peptide side chains of peptidoglycan (PG) strands during biosynthesis of the PG network, which builds the backbone of the cell wall. The *pbpB1* gene (*lmo1438*) of *L. monocytogenes* is essential (10) and depletion of PBP B1 prevents growth (9). Moreover, PBP B1 is required for maintenance of the rod-shaped cellular morphology and resistance against (amino)penicillins (9), reflecting the importance of PBP B1 for PG biosynthesis and viability as well as its relevance as a drug target.

Next to (amino)penicillins, carbapenems are also occasionally used in the therapy of listeriosis, but treatment failure (11) and appearance of spontaneously occurring resistant suppressors have been reported (8, 12). Interestingly, imipenem resistant suppressors isolated during *in vitro* culture in the presence of imipenem altered PBP B1 in a way that reduced its affinity for penicillin causing resistance (8, 12). The reason for this effect was not yet known at this time, but mutations in *pbpB1* itself were most likely responsible.

Resistance against (amino)penicillins has occasionally been detected in food and environmental isolates of *L. monocytogenes* (*13*), whereas β-lactam-resistant clinical strains have been described less frequently (14, 15) and were even absent in recent nationwide screenings of large isolate collections for phenotypic antimicrobial resistance (16, 17). Because the few β-lactam-resistant isolates identified have never been investigated further, the molecular basis of β-lactam resistance in *L. monocytogenes* is unknown.

In this work, we describe the detection and further characterization of an ampicillin resistant *L. monocytogenes* isolate obtained from a listeriosis patient in Germany in 2023. Whole genome sequencing and genetic follow-up experiments showed that a specific *pbpB1* mutation caused ampicillin resistance in this strain. The same mutation was acquired during ampicillin selection *in vitro*. Moreover, it also was found in other clinical isolates, which then turned out to be ampicillin resistant.

## Materials and Methods

### Bacterial strains and growth conditions

Clinical *L. monocytogenes* strains were isolated from infected patients in primary diagnostic labs and sent to the Consultant laboratory for *Listeria* at unit 11 of the Robert Koch Institute. Tab. 1 lists all bacterial strains and plasmids used in this study. *L. monocytogenes* strains were generally grown in BHI broth or on BHI agar plates at 37°C. Erythromycin (5 µg ml^-1^) and X-Gal (100 µg ml^-1^) were added where necessary. *Escherichia coli* TOP10 was used as the standard cloning host.

**Table 1:**
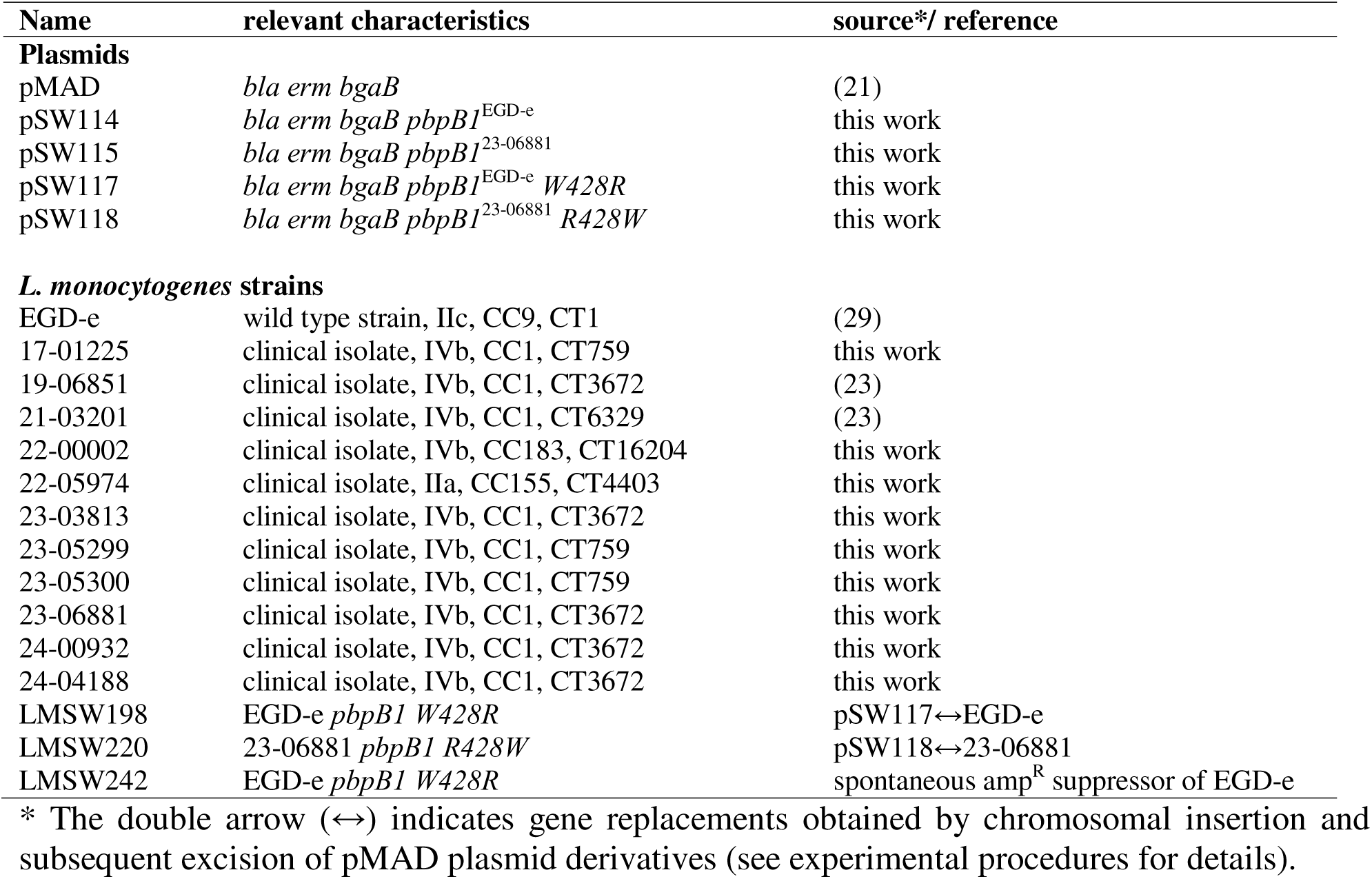
Plasmids and strains used in this study.

### General methods, manipulation of DNA and oligonucleotide primers

Standard methods were used for transformation of *E. coli* and for isolation of plasmid DNA. Transformation of *L. monocytogenes* was performed using electroporation as described by others (18). PCR, restriction and ligation of DNA was performed following the manufactureŕs instructions. All primer sequences are listed in Tab. 2.

**Table 2:**
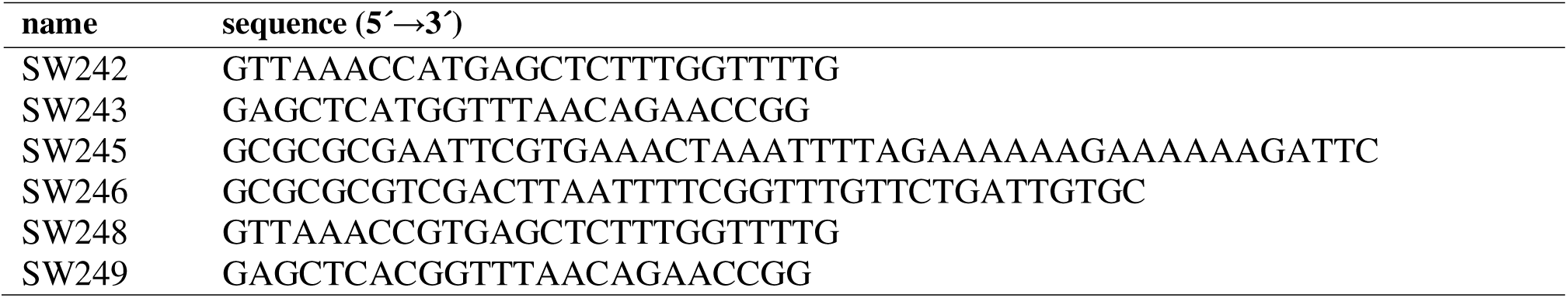
Primers used in this study.

### Whole genome sequencing, *in silico* serogrouping, MLST, cgMLST and agMLST

Isolation of genomic DNA was performed by mechanical disruption using glass beads in a TissueLyser II bead mill (19). Libraries were prepared using the Nextera XT DNA Library Prep Kit and sequenced on MiSeq or NextSeq sequencers in 2 x 250 bp or 2 x 150 bp paired end mode, respectively. Read trimming and assembly with Velvet as the assembler was performed in SeqSphere+ v10 (Ridom, Germany). *In silico* molecular serogroups, seven locus multi-locus sequence typing (MLST) sequence types (STs), 1,701 locus core genome MLST (cgMLST) complex types (CTs) and the 1,158 MLST alleles of the accessory genome (agMLST) were automatically extracted using SeqSphere+ (20). cgMLST clusters and minimum spanning trees were calculated in SeqSphere+ in the “pairwise ignore missing values” mode. Sequence alignments were also generated in SeqSphere+. All genome sequencing data are available at the <u>European Nucleotide Archive</u> using the accession numbers given in supplementary Table S1.

### Construction of plasmids and strains

The *pbpB1* gene of *L. monocytogenes* strain EGD-e (*lmo1438*) was amplified from chromosomal DNA in a PCR using the oligonucleotides SW245/SW246 as the primers. The obtained *pbpB1* fragment was inserted into plasmid pMAD using EcoRI/SalI as the restriction enzymes. The W428R mutation was then introduced into the *pbpB1* gene of the resulting plasmid (pSW114) by quikchange mutagenesis using the oligonucleotides SW242/SW243 as the mutagenic primers, yielding plasmid pSW117. Likewise, *pbpB1* of strain 23-06681 was amplified by PCR using SW245/SW246 as the primers and cloned into pMAD after EcoRI/SalI restriction, which yielded plasmid pSW115. The W428R mutation present in this plasmid was then corrected by quikchange mutagenesis using SW248/SW249 as the primers.

Both plasmids were introduced into the recipient strains by electroporation (18) and erythromycin resistant clones were selected. Allelic exchange was performed following the protocol of Arnaud and colleagues (21). Introduction of the desired sequence changes into *pbpB1* was confirmed by genome sequencing.

### Determination of minimal inhibitory concentrations

Ampicillin, penicillin G and meropenem were purchased from Sigma-Aldrich. For the determination of minimal inhibitory concentrations (MIC), an overnight culture was diluted to an OD_600_ of 0.1 in BHI broth. Next, a 96 well cell culture plate was loaded with 100 µl BHI medium containing twice the concentration of the antibiotic to be tested as a two-fold dilution series. 100 µl of the diluted overnight culture was added to each well and growth was recorded at 37°C for 24 h in an automated fashion using a Multiskan Sky or Multiskan Go Microplate Spectrophotometer (Thermo Fisher Scientific). The MIC was defined as the lowest concentration where no bacterial growth was observed after 20 hours. Where available, clinical breakpoints for *L. monocytogenes* provided by the EUCAST were used for interpretation of MIC values (22).

## Results

### An ampicillin resistant *L. monocytogenes* CC1 strain from a German listeriosis patient

The German consultant laboratory for *Listeria* (CL) at the Robert Koch Institute collects 500-600 *L. monocytogenes* isolates per year from listeriosis patients in Germany for genomic surveillance (23). In October 2023, a clinical *L. monocytogenes* isolate was received from a primary diagnostic laboratory with suspected ampicillin resistance. This isolate (internal identification number: 23-06881) had been collected from a 77-year old female patient suffering from an exacerbating chronic obstructive pulmonary disease. The patient had received cortisone therapy and piperacillin/tazobactam treatment, which was later switched to meropenem, due to rising clinical infection parameters of unknown cause. *L. monocytogenes* was isolated from one blood culture and antibiotic susceptibility testing of the obtained isolate indicated ampicillin resistance. Although the antibiotic treatment was changed to cotrimoxazole, the infection was fatal. The isolate 23-06881 belonged to serogroup IVb and it was subtyped as CC1 (ST698) by MLST and as CT3672 by cgMLST after whole genome sequencing. Furthermore, it belonged to a small cluster consisting of eight genetically closely related CC1 isolates (internal RKI cluster name: Tau2a, Fig. 1A), which had been collected from eight listeriosis patients in Germany between 2019-2024. Determination of the ampicillin MIC of isolate 23-06881, of several isolates representing the diversity within the Tau2a cluster and of a related non-Tau2a CC1 isolate from another recent outbreak (21-03201, cluster Alpha10) (23) confirmed ampicillin resistance of isolate 23-06881 (MIC: 2±0 µg/ml), while all other tested strains were susceptible (Fig. 1B). This demonstrates that the occurrence of ampicillin resistance in this phylogenetic branch of the *L. monocytogenes* population must have been a sporadic event.

**Figure 1:**
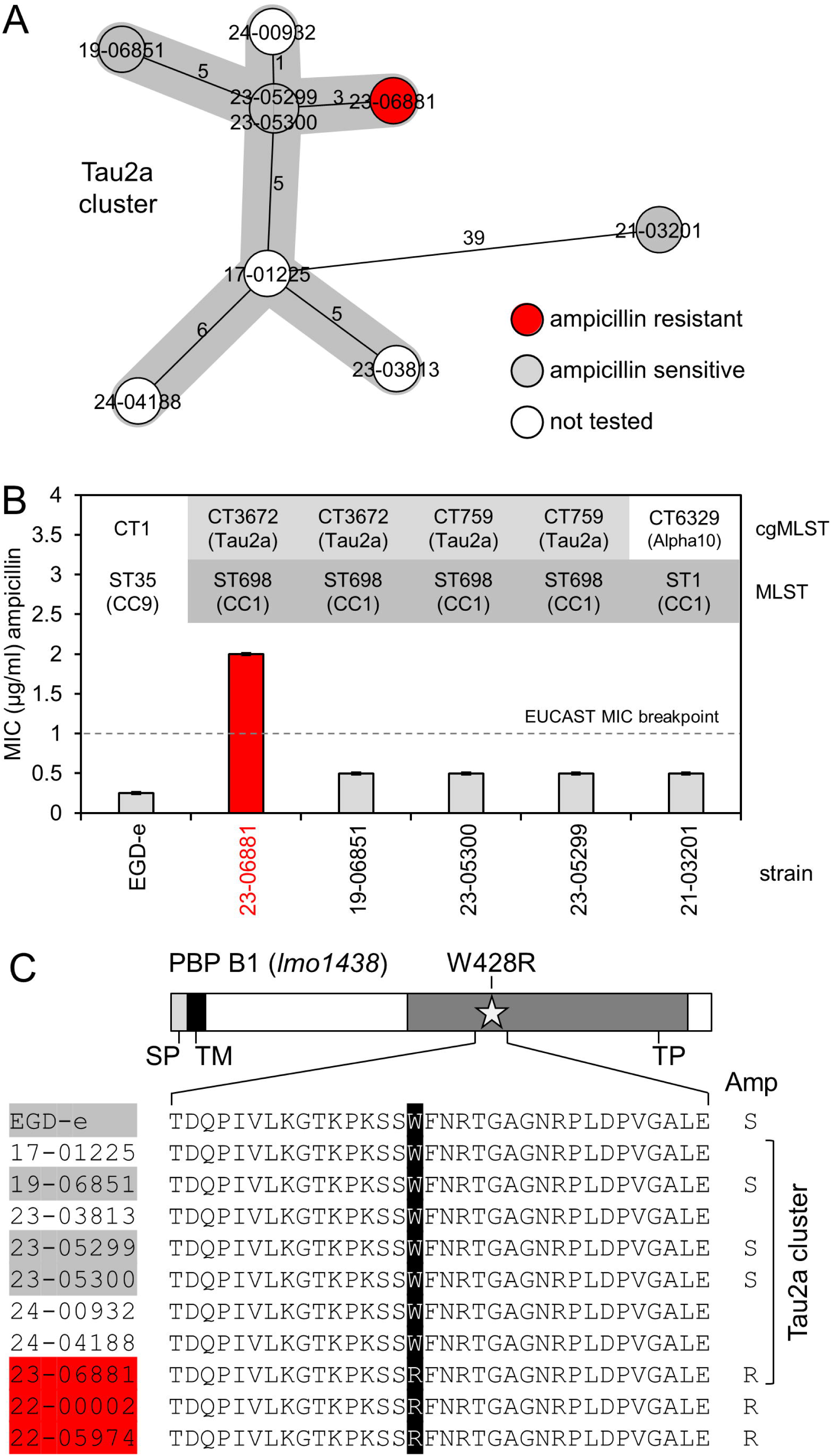
An ampicillin resistant *L. monocytogenes* CC1 strain with a *pbpB1* mutation. (A) Minimal spanning showing the phylogenetic CC1 cluster (named “Tau2a”), in which ampicillin resistance was discovered. The CC1 strain 21-03201 (a representative of another CC1 cluster, called Alpha1) was included as outgroup. The tree was calculated from 1701 locus cgMLST data and strains are coloured according to their ampicillin resistance or sensitivity. (B) Minimal inhibitory concentrations (MICs) of ampicillin of a subset of strains shown in panel A. *L. monocytogenes* reference strain EGD-e was included as control. Standard deviations are calculated from MICs, which have been determined three independent times. cgMLST and MLST typing data are included. (C) The ampicillin resistant *L. monocytogenes* Tau2a strain 23-06881 carries an amino acid exchange in the *pbpB1* gene. Upper part: Scheme showing the domain organization of PBP B1 (abbreviations: SP – signal peptide, TM – transmembrane segment, TP – transpeptidase domain). Lower part: Section of a multiple sequence alignment of the *pbpB1* gene of the *L. monocytogenes* strains belonging to the Tau2a cluster shown in panel A and of strains 22-0002 and 22-05974 also carrying the W428R mutation. The W428R mutations is highlighted and strain names are coloured according to their experimentally determined ampicillin resistance. S – sensitive, R – resistant.

### The ampicillin resistant isolate 23-06881 carries a point mutation in *pbpB1*

In order to identify the genetic basis of this resistance phenotype, we first searched the genome of strain 23-06881 for the presence of reads specific for any of the genes belonging to the six *Listeria repA*-family plasmids (24), but evidence for the presence of a plasmid was not found. Therefore, we compared the genome of strain 23-06881 with the genomes of the seven other isolates of the Tau2a cluster. For this, we searched for cgMLST and agMLST alleles that were specific to 23-06881 using the group-specific single nucleotide variant search tool in SeqSphere, which identified two non-synonymous SNPs that were specific to isolate 23-06881: A W428R substitution in the *pbpB1* gene (*lmo1438*) encoding penicillin binding protein (PBP) B1 as well as a N276K substitution in *purB* (*lmo1773*) that encodes adenylosuccinate lyase. While the latter enzyme is required for purine biosynthesis, particularly during intracellular growth (10), PBP B1 is an essential transpeptidase mediating crosslinking of peptidoglycan (PG) strands during cell wall biosynthesis (9, 10). PBP B1 was linked previously to ampicillin resistance in the *L. monocytogenes* reference strain EGD-e (9) and therefore we reasoned that the *pbpB1* W428R mutation most likely would explain ampicillin resistance in isolate 23-06881.

### Transplantation and correction of the *pbpB1 W428R* mutation affects β-lactam resistance

To test the effect of the *pbpB1 W428R* mutation on β-lactam resistance, the W428R substitution was transplanted to the *pbpB1* gene of the ampicillin susceptible *L. monocytogenes* reference strain EGD-e. The MIC of ampicillin of the resulting strain (LMSW198) was four times higher (1.0±0 µg/ml) than that of strain EGD-e (0.25±0 µg/ml µg/ml). This is not considered clinical resistance yet, as the MIC was just as high but not higher than the current EUCAST MIC breakpoint for ampicillin (Fig. 2A) (22). Likewise, resistance to amoxicillin was increased fourfold (Fig. 2A). In contrast, resistance against penicillin G was not affected by the *pbpB1 W428R* mutation (Fig. 2A). However, transplantation of the W428R mutation rendered strain EGD-e, which is normally sensitive to meropenem (MIC: 0.09±0.03 µg/ml), fully meropenem resistant (MIC: 3±1 µg/ml). To further confirm these findings, the W428R substitution in the *pbpB1* gene of the ampicillin resistant clinical isolate 23-06881 was corrected to tryptophan. This reduced resistance to ampicillin and amoxicillin four- and eightfold, respectively, but it did not affect penicillin resistance. However, meropenem resistance of isolate 23-06881 (MIC: 6±2 µg/ml) was lost upon correction of the W428R exchange to wildtype tryptophan (MIC: 0.2±0.06 µg/ml, Fig. 2A). These experiments demonstrate that the W428R mutation present in the *pbpB1* gene of strain 23-06881 was the sole reason for its resistance against ampicillin, amoxicillin and meropenem.

**Figure 2:**
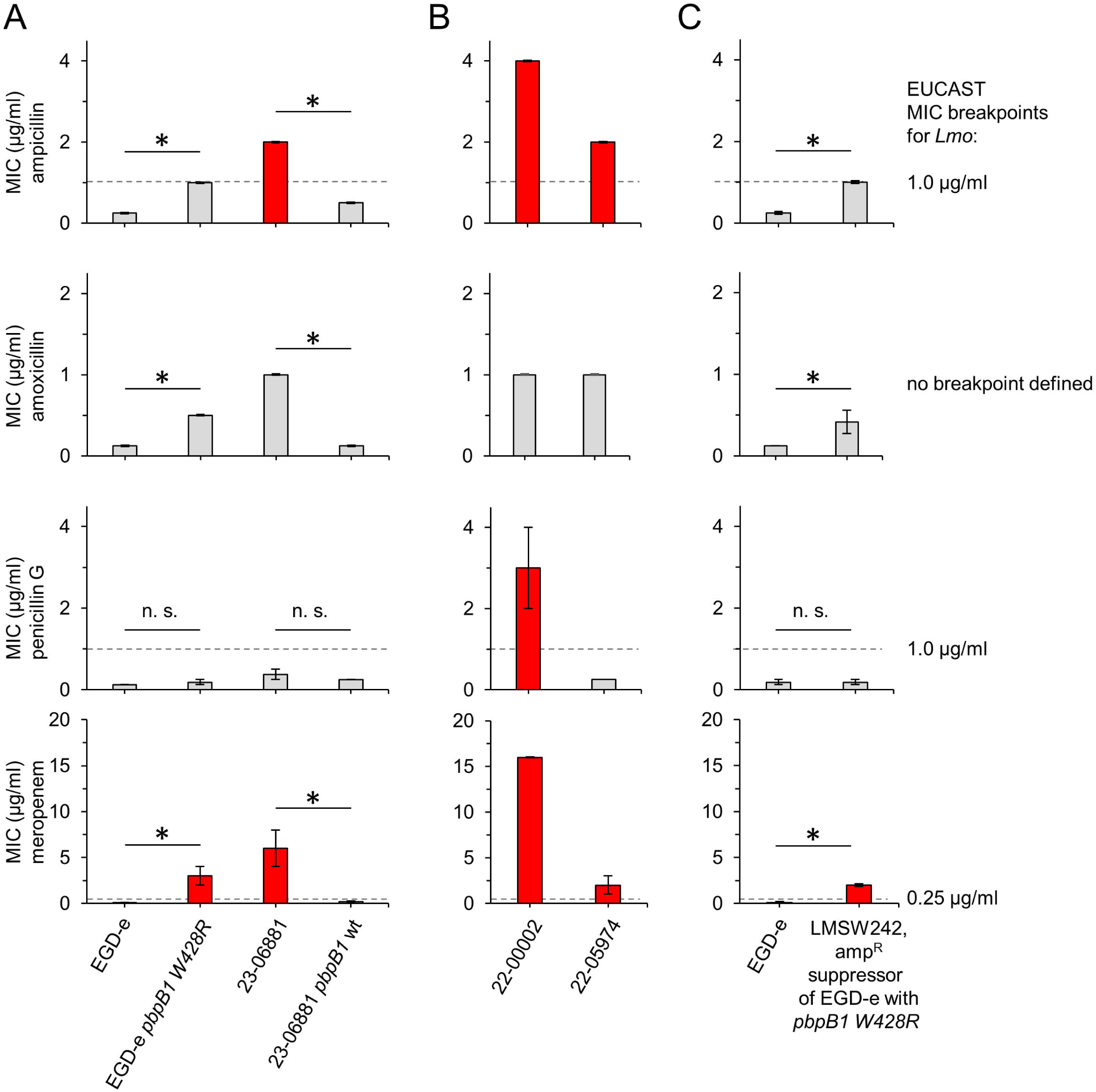
The *pbpB1 W428R* mutation causes *L. monocytogenes* β-lactam resistance. (A) Introduction of the *pbpB1 W428R* mutation into strain EGD-e and correction of this mutation in the ampicillin resistant 23-06881 background. MICs of ampicillin, amoxicillin, penicillin G and meropenem for *L. monocytogenes* strains EGD-e, LMSW198 (EGD-e *pbpB1 W428R*), 23-06881 and LMSW220 (23-06881 *pbpB1* wt) are shown. (B) Resistance of further clinical *L. monocytogenes* isolates carrying the *pbpB1 W428R* mutation against the same set of β-lactams. (C) β-lactam resistance of a spontaneous *pbpB1 W428R* mutant selected *in vitro*. Minimal inhibitory ampicillin, amoxicillin, penicillin G and meropenem concentrations for *L. monocytogenes* strains EGD-e (wild type reference strain) and LMSW242. Strain LMSW242 is a descendant of EGD-e and has spontaneously acquired the *pbpB1 W428R* mutation during *in vitro* selection in the presence of ampicillin. MICs were determined three independent times and average values and standard deviations were calculated. Asterisks indicate significance levels *(P*<0.05, n=3). Values that exceed the EUCAST breakpoints are colored red.

### Occurrence of the *pbpB1 W428R* mutation in other clinical *L. monocytogenes* isolates of different phylogenetic sublineages

We searched our collection of *L. monocytogenes* isolates for the presence of other isolates carrying the *pbpB1 W428R* mutation. This collection comprised approximately 4300 genome-sequenced isolates from listeriosis patients in Germany from 2007-2024 at the time of analysis. Extraction of *pbpB1* sequences and their inspection resulted in the identification of two more recent clinical isolates also carrying the same mutation (Fig. 1C): The first isolate (internal ID: 22-00002) is a sporadic serogroup IVb-v1 isolate belonging to clonal complex CC183 (cgMLST complex type CT16204) and was collected from a sepsis patient with reported aneurysm. This isolate had an even higher resistance against ampicillin (4±0 µg/ml), penicillin (3±1 µg/ml) and meropenem (16±0 µg/ml) than the originally identified 23-06881 isolate (Fig. 2B). Thus, additional genetic markers in addition to *pbpB1 W428R* further raising β-lactam resistance must be present. The second isolate (22–05974) is a sporadic serogroup IIa isolate belonging to CC155 (CT4403) and was collected from a tissue specimen of another patient with reported aneurysm. Ampicillin, amoxicillin, penicillin and meropenem resistance levels of this second isolate were comparable to that of strain 23-06881 (Fig. 2B).

### Spontaneous aquisition of the *pbpB1 W428R* mutation by ampicillin selection *in vitro*

Next, we wondered whether the *pbpB1 W428R* mutation could result from genomic adaptation to increased ampicillin levels and hence cultivated *L. monocytogenes* strain EGD-e in the presence of an increased ampicillin concentration (twice the MIC of EGD-e) to select for spontaneously arising ampicillin resistant suppressors. We in fact observed occasional initiation of growth approximately 20 hours after inoculation. Two potential suppressors were isolated from these cultures and their genomes sequenced. Both carried the *pbpB1 W428R* mutation as the sole genetic change. The MICs for ampicillin, amoxicillin, penicillin and meropenem were determined for one of them (LMSW242), which showed that the MIC of ampicillin was increased four-fold, of amoxicillin three-fold and that of meropenem 32-fold, whereas the MIC of penicillin was not affected (Fig. 2C). Thus, this strain behaves identical as strain LMSW198, which was generated from EGD-e by introduction of the *pbpB1 W428R* mutation using homologous recombination techniques (see Fig. 2A).

Strains carrying the *pbpB1 W428R* mutation did not show a growth defect (Fig. 3A), despite *pbpB1* essentiality (9). Since this mutation is only infrequently observed, we wondered as to when this mutation would be disadvantageous and studied growth of the *pbpB1 W428R* strain under several conditions. We observed that the *pbpB1 W428R* mutation impairs growth at 42°C (Fig. 3A) and decreases resistance of *L. monocytogenes* to ceftriaxone (Fig. 3B). *L. monocytogenes* is naturally resistant to cephalosporins such as ceftriaxone, suggesting that constant selection for maintenance of full cephalosporin resistance could suppresses acquisition of the *pbpB1 W428R* mutation in its natural reservoirs.

**Figure 3:**
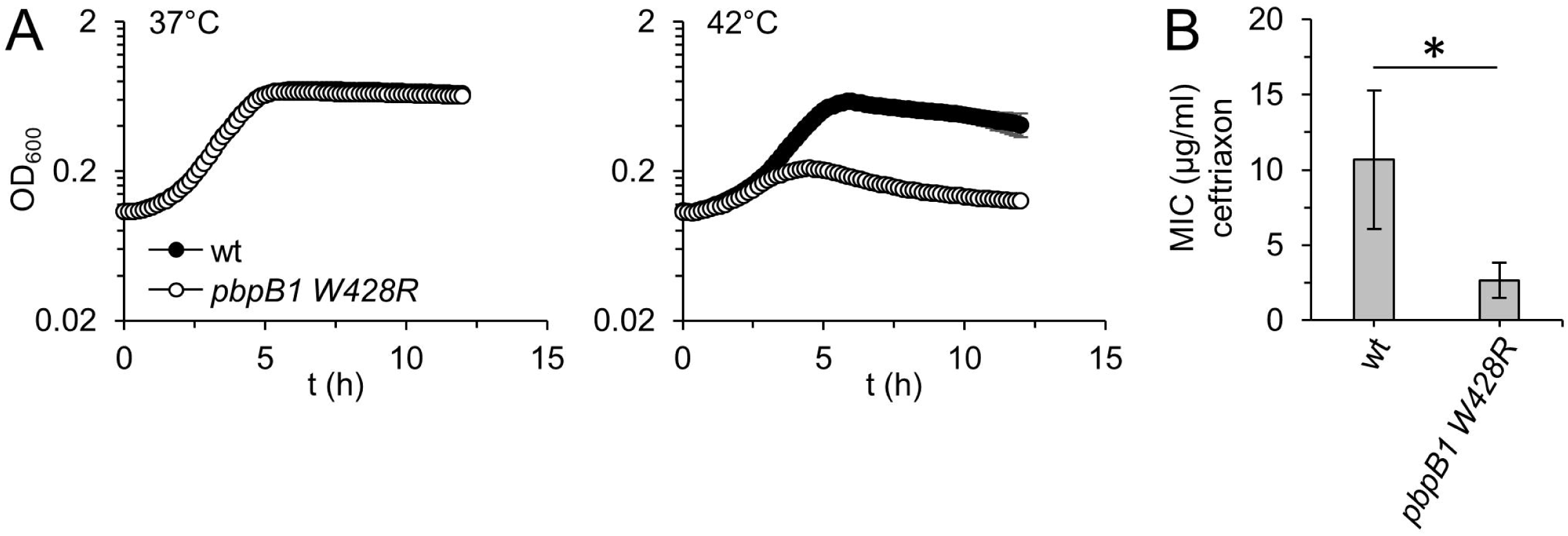
Detrimental effects of the *pbpB1 W428R* mutation. (A) Growth of *L. monocytogenes* strains EGD-e (wt) and LMSW198 (*pbpB1 W428R*) in BHI broth at 37°C (left) or 42°C (right). Growth curves are average values calculated from technical triplicates and standard deviations were included. (B) Minimal inhibitory concentration of ceftriaxone for *L. monocytogenes* strains EGD-e (wt) and LMSW198 (*pbpB1 W428R*). MICs were determined three independent times and average values and standard deviations were calculated. Asterisks indicate significance levels *(P*<0.05, n=3).

## Discussion

Here we describe clinical *L. monocytogenes* isolates that are resistant to ampicillin and the identification of a mutation in the *pbpB1* gene as cause for this resistance phenotype. Trp-428 of PBP B1 was exchanged for Arg in these isolates, whereby the bulky heterocyclic side chain of tryptophan was replaced by the positively charged side chain of arginine. Trp-428 is located in the transpeptidase domain of PBP B1 at the tip of a loop that lies between the conserved SxxK and SxN motifs that are both part of the active site (25). This loop folds in the direction of the active site pocket generating a distance of only ∼10 Å between Trp-428 and the active site serine in the Alphafold crystal structure model of PBP B1 (Fig. 4). It seems plausible that introduction of additional positive charge at this position may affect accessibility of the active site for inhibitors, such as ampicillin. In good agreement, homologs of PBP B1 in other *Bacillota* such as *Staphylococcus aureus* PBP3 or *Streptococcus pneumoniae* PBP2b have also been linked to β-lactam resistance when specific amino acid substitutions occurred in their transpeptidase domains (26, 27), although at different sites.

**Figure 4:**
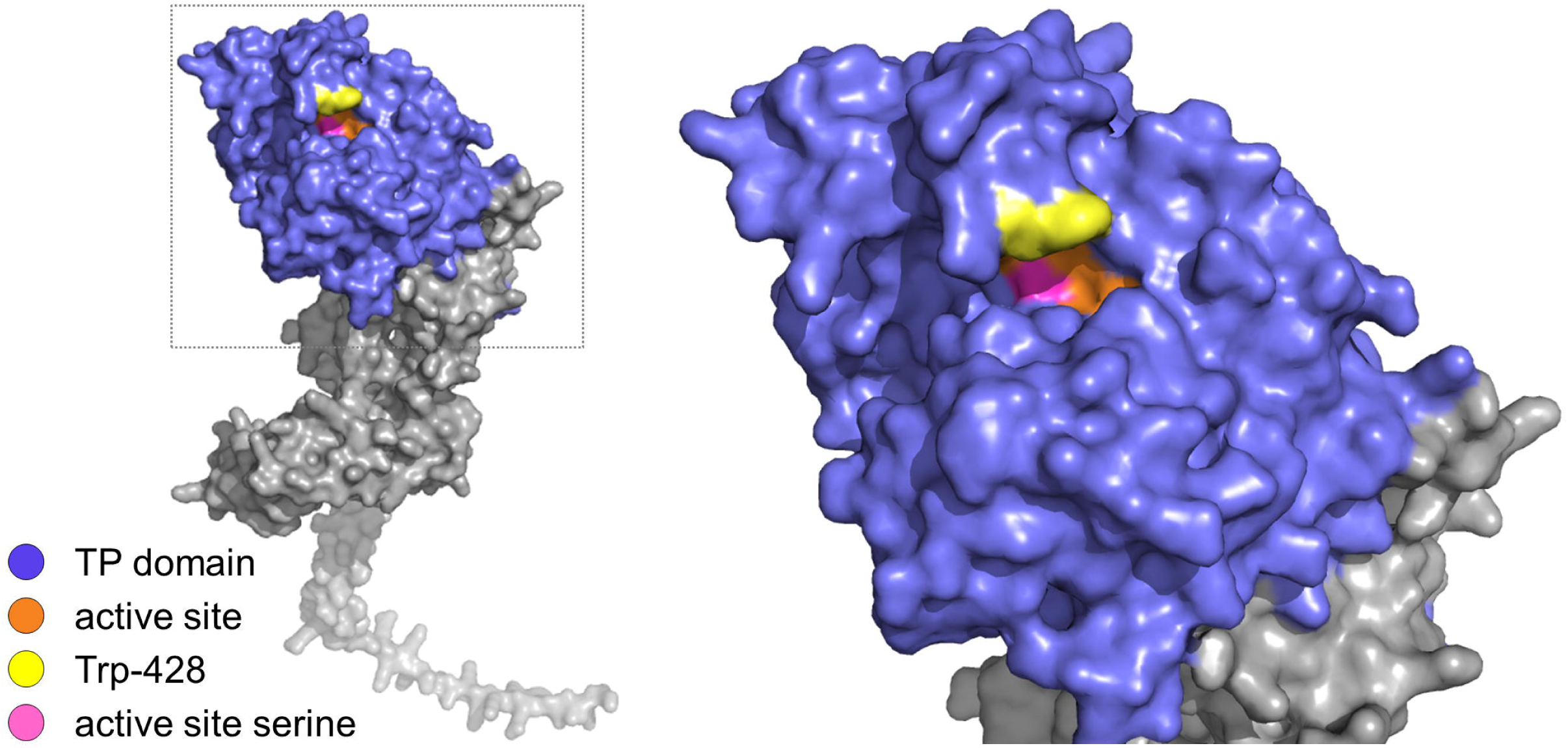
Trp-428 is located in the close vicinity of the active site of PBP B1. Alphafold model of *L. monocytogenes* PBP B1 (30). The transpeptidase (TP) domain is highlighted in the model of the full-length protein (left) and shown separately at higher magnification (right). The active site, the catalytic serine and the position of the Trp-428 residue, which is mutated in the ampicillin-resistant *L. monocytogenes* isolates, are also highlighted.

Ampicillin resistance either occurred in sporadic isolates (22–00002, 22–05974) or only once among the otherwise genetically closely related isolates of the same outbreak cluster (23–06881). In addition, all three ampicillin-resistant isolates identified here were representatives of different molecular serogroups and phylogenetic sublineages. This indicates that ampicillin resistance due to *pbpB1* mutation is not a monophyletic characteristic of a specific phylogenetic sub-lineage, but instead must be seen as the result of independent mutational events. All three patients had a common history of piperacillin/tazobactam treatment (followed by meropenem administration in two out of the three cases) prior to diagnosis of *L. monocytogenes*. Piperacillin/tazobactam is not occurred in the infected patients during antibiosis. In good agreement with this, acquisition of the *pbpB1 W428R* mutation was detected *in vitro* during ampicillin selection *in vitro*, where the appearance of resistant suppressors was frequently observed. Since *L. monocytogenes* isolates are usually sent to the CL immediately after diagnosis, we assume that selection of resistant suppressors occurs more frequently during antibiotic treatment than we can determine by systematic characterization of submitted isolates. With penicillin, however, we have never observed the occurrence of resistant suppressors in our selection experiments with strain EGD-e. The *pbpB1 W428R* mutation was detected with a frequency of about 10^-3^ in the strain collection of the CL. However, there must be further mutations that can raise β-lactam resistance to an even higher level than originally observed with the ampicillin-resistant 23-06881 isolate, since isolate 22-00002 is approximately twice as resistant against ampicillin and meropenem and reaches complete penicillin resistance, for which the presence of the *pbpB1 W428R* mutation was not sufficient (Fig. 2B). Comparison of the genome of penicillin-resistant isolate 22-00002 with genomes of penicillin-sensitive isolates of the same sub-lineage (CC183) (16) identified 19 non-synonymous SNPs that are possibly involved, but whose individual contribution to penicillin resistance was not immediately obvious. Nevertheless, this shows that further mutations likely exist that also promote resistance of *L. monocytogenes* against penicillin so that we have to assume that the frequency of *L. monocytogenes* isolates resistant against these antibiotics is presumably somewhat higher than estimated above. Continuous screening of clinical *L. monocytogenes* isolates for resistance against penicillins is therefore justified, especially when a history of prolonged antibiotic treatment has been documented.

Lastly, it is important to point out that isolate 23-06681, where the *pbpB1 W428R* mutation was initially discovered, belongs to the CC1 sub-lineage. Infections with CC1 isolates are associated with an increased risk for the development of neurolisteriosis and pregnancy-associated listeriosis (23, 28). Thus, our work shows that ampicillin resistance can develop in hypervirulent sub-lineages of *L. monocytogenes*. Moreover, it indicates that antibiotic susceptibility testing should be part of standard listeriosis therapy.

## Acknowledgments

We thank Simone Dumschat for excellent technical assistance and the Genome Competence Centre of the RKI for sequencing of *L. monocytogenes* genomes.

